# The SARS-CoV-2 nucleocapsid protein inhibits the cellular Nonsense-Mediated mRNA Decay (NMD) pathway preventing the full enzymatic activation of UPF1

**DOI:** 10.1101/2024.07.16.603692

**Authors:** Veronica Nuccetelli, Makram Mghezzi-Habellah, Séverine Deymier, Armelle Roisin, Francine Gerard-Baraggia, Cecilia Rocchi, Pierre-Damien Coureux, Patrice Gouet, Andrea Cimarelli, Vincent Mocquet, Francesca Fiorini

## Abstract

The Nonsense-mediated mRNA decay (NMD) pathway triggers the degradation of defective mRNAs and governs the expression of mRNAs with specific characteristics. Current understanding indicates that NMD is often significantly suppressed during viral infections to protect the viral genome. In numerous viruses, this inhibition is achieved through direct or indirect interference with the RNA helicase UPF1, thereby promoting viral replication and enhancing pathogenesis.

In this study, we employed biochemical, biophysical assays, and cellular investigations to explore the interplay between UPF1 and the Nucleocapsid (Np) protein of SARS-CoV-2. We evaluated their direct interaction and its impact on inhibiting cellular NMD. Furthermore, we characterized how this interaction affects UPF1’s enzymatic function. Our findings demonstrate that Np inhibits the unwinding activity of UPF1 by physically obstructing its access to structured nucleic acid substrates. Additionally, we showed that Np binds directly to UPF2, disrupting the formation of the UPF1/UPF2 complex essential for NMD progression. Intriguingly, our research also uncovered a surprising pro-viral role of UPF1 and an antiviral function of UPF2.

These results unveil a novel, multi-faceted mechanism by which SARS-CoV-2 evades the host’s defenses and manipulates cellular components. This underscores the potential therapeutic strategy of targeting Np-UPF1/UPF2 interactions to treat COVID-19.

## Introduction

Nonsense-mediated mRNA decay (NMD) was originally described as a eukaryotic translation-dependent quality control mechanism that degrades mRNAs harboring premature translation termination codons (PTC; (1,2)). The NMD exerts a broader global influence by regulating the stability of numerous error-free transcripts, thereby influencing 5-20% of cellular gene expression (3). Beyond the PTC, all the NMD-prone transcripts possess unique characteristics, such as the long 3’ untranslated regions (UTRs), exon junction complexes (EJCs) in the 3’UTR or a short open reading frames in the 5’UTR (uORFs), which make them susceptible to NMD (4). Recent studies have unveiled the role of NMD as restrictor of RNA virus replication (reviewed in (5–7). This phenomenon is likely attributed to the replication strategy of RNA- and retro-viruses, which undergo selective pressure to maximize the coding capacity of their viral RNA without increasing the size of their genome. As a result, certain features such as internal termination codons (iTC) due to multiple ORFs and long 3’ UTRs, are introduced into viral RNAs and make them prone to NMD. The list of phyto- and mammalian viruses sensitive to NMD restriction continues to expand, encompassing numerous families (8–26). These viruses have evolved by developing strategies to counteract NMD and sometime co-opting the NMD factors for their advantage. They employ either RNA structures that directly *cis*-protects the viral genome (9,10,27–29) or *trans*-acting factors that target various NMD proteins (8,13–20,23–25).

The RNA helicase UPF1 (UP-frameshift 1) is the central factor of NMD. This multifunctional enzyme comprises a conserved helicase core domain that, powered by ATP hydrolysis, is capable of unwinding double-stranded nucleic acids in a 5’ to 3’ direction. UPF1 is a dynamic nucleoprotein complexes remodeler: it moves along single-stranded nucleic acids displacing the proteins that are tightly attached to the substrate. The activity of UPF1 is inhibited by two domains that surround the helicase core: the N-terminal CH domain, which is rich in cysteine and histidine residues, and the C-terminal SQ domain, which contains clusters of serine-glutamine residues. To date, the only enzymatic activation of UPF1 that has been characterized during NMD is that exerted by UPF2. Indeed, the binding of UPF2 to the CH domain allows for its displacement, inducing the activation of UPF1 by shifting the enzyme from an RNA-clamping to a relaxed-translocating mode. Few viral factors have been described to integrate this complex enzymatic regulation and affect NMD. For example, the Tax protein of Human T-lymphotropic virus 1 retrovirus (HTLV-1) directly binds to both UPF1 and UPF2 (11). Tax, by interacting with the helicase core domain of UPF1, induces a loss of affinity for the substrate resulting in blocking of ATPase and translocation activities (19). It has also been reported that HTLV-1 Rex inhibits NMD via interaction with UPF1 but molecular details are not yet available (25,30). An interesting case is represented by human immunodeficiency virus-1 (HIV-1) that not only evades NMD but usurps UPF1 to promote viral replication (14,31,32). In this context, UPF1 has a pro-viral effect by contributing to enhancing the viral RNA stability and translation and the assembly of new virions (14,31–33). This pro-viral function of UPF1 is independent of UPF2, which is excluded from UPF1-containing viral RNPs. UPF2 has a genuine antiviral activity by inhibiting HIV-1 replication by promoting viral RNA decay (14,33).

RNA viruses of the Flaviviridae family also disrupt NMD through various mechanisms (17,20,21). Among them, Zika virus (ZIKV) directly targets UPF1 via its capsid (CA): the protein which interacts with and packages viral RNA (Fontaine 2018). This interaction occurs in both the nucleus and cytoplasm and appears to lead to the specific proteasomal degradation of nuclear UPF1 (17). The implications of this nuclear interaction for NMD and the effect of ZIKV CA on the activity of cytoplasmic UPF1 need further clarification.

The interplay between NMD and the Mouse Hepatitis Virus (MHV), a prototypic member of the Betacoronavirus (β-CoV) gender to which belong SARS (Severe Acute Respiratory Syndrome) and MERS (Middle East respiratory Syndrome) viruses, is well described (18). In this study, the authors showed that MHV genomic and some sub-genomic RNAs are NMD targets. Indeed, depletion of NMD factors such as UPF1 and UPF2, promote the accumulation of viral RNAs early in infection, leading to the production of higher titers of MHV and enhanced replication (16). The MHV factor that antagonizes NMD is the nucleocapsid protein (Np), a viral structural protein that binds to and organizes the viral genome within virions (18,34). Consistently with this results, two proteomic studies identified UPF1 as a potential molecular partner of Np in Infectious Bronchitis Virus (IBV) Coronavirus and SARS-CoV-2 (35,36). Notably, UPF1 has also been shown to directly and specifically bind to SARS-CoV-2 genomic and sub-genomic RNAs in infected human cells by two research groups (37,38). However, the direct interaction between CoV Np and UPF has not yet been confirmed, and mechanistic insights into NMD inhibition are also unknown.

In this study, we used biochemical and biophysical assays combined with cellular studies to understand whether and how SARS-CoV-2 manages the NMD pathway. We elucidated the molecular mechanism through which the SARS-CoV-2 Np protein impacts the activity of UPF1, leading to the inhibition of cellular NMD. Np exerts a dual-control mechanism on UPF1 by inhibiting both its activity and its activation and hijacking it for its advantage. Our study provides valuable molecular and cellular insights into this intricate inhibitory mechanism and underscores the significance of NMD in restricting SARS-CoV-2.

## Materials and Methods

### Cell culture

Vero E6, HEK 293T and HeLa cells were cultured in DMEM (invitrogen) complemented with 5% FCS and penicillin/streptomycin at 37°C, 5%CO2.

### NMD assay

NMD assays were performed as described previously (19): 0.5 μg of a β-globin reporter minigene either with a WT sequence (Gl-WT) or with a PTC in the second exon (Gl-PTC) was transfected in 0.6×106 HeLa cells with jetprime reagent (polyplus transfection). When indicated, 0.5 µg of the SARS-CoV-2 Np coding plasmid (Addgene, cat.141391) was co-transfected as well. In the case of UPF1 extinction experiments, siRNA duplexes (either control (mission siRNA universal negative control, SIGMA) or targeting UPF1 (5-GAUGCAGUUCCGCUCCAUU-3) were transfected alongside with the plasmid following the manufacturer recommendations. The medium was changed after 12 h, and cells were further cultivated for 24 additional hours. Total mRNAs were extracted using the Macherey-Nagel RNA easy extraction kit and quantified by RT-qPCR using the QuantiTect SYBR Green qRTPCR kit (Qiagen) and the following primers (5-TTGGGGATCTGTCCACTCC-3_, 5-CACACCAGCCACCACTTTC-3). Normalisation was carried out with respect to renilla mRNA (primers: 5_-CTAACCTCGCCCTTCTCCTT-3, 5_-TCGTCCATGCTGAGAGTGTC-3) expressed from a cotransfected Renilla luciferase plasmid (0.2µg) and NMD insensitive. The values represented in the graphs correspond to the mean of at least three biological replicates, and the error bars correspond to the SD. Half-lives were calculated for each replicate, and P values were calculated by performing a Student’s t-test (unpaired, two-tailed) ns: P > 0.05; *P < 0.05.

### Viral replication

We used a replicative SARS-CoV-2 (Wuhan strain) bearing the mNeongreen reporter replacing the ORF7, kindly obtained from Pei-Yong Shi at the University of Texas Medical Branch, Galveston, TX, USA (39). SARS-CoV-2 mNeonGreen viral stocks were produced in Vero E6 cells. For infection experiments, Vero E6 cells were transfected with 4 µg of DNAs coding for HA-UPF1, HA-UPF2 or a pCMV empty as a negative control with jet prime reagent. Cells were challenged 28 hours later with SARS-CoV-2 mNeonGreen at multiplicities of infection (MOIs) comprised between 0,1 and 0,01 and the extent of viral replication was determined 18 hours post infection by flow cytometry on a MacsQuant cytometer.

### Protein expression and purification

Human UPF1-HD (295-914 aa), UPF1-CH-HD (115-914 aa) and UPF2 (704–1204) proteins were expressed and purified as described in (40,41). SARS-CoV-2 Np protein fragments (Np-L (50-370aa), Np-NTD (50-138 aa), Np-CTD (250-370 aa); Np-NTD-SRD (50-250 aa)) were cloned in pET52b+ in frame with a C-terminal 10xHistidine tag by standard molecular biology techniques. Recombinant proteins were produced from E.coli BL21(DE3) Rosetta bacteria (Sigma Aldrich, cat. 71403). Bacterial cells were grown in an LB medium for 16 hours at 25°C, then protein expression was induced with 0.1 mM IPTG (isopropyl-β-dthiogalactopyranoside, Sigma Aldrich, cat. PHG000110). Cultures were incubated 6h at 18°C under continuous shaking then harvested by centrifugation for 15 minutes at 5000 g. Each pellet was resuspended with buffer A (50 mM Tris-HCl pH 7.5, 150 mM NaCl, 20 mM imidazole, 5 mM β-mercaptoethanol, 0.5 mM CaCl2, and 20% glycerol, v/v) supplemented with 1 mM of adenosine triphosphate (ATP, Jena Bioscience cat. NU-1010), 1 mM MgCl2, DNase I (New England Biolabs, Cat. M0303S), RNaseA (Sigma Aldrich Cat. RNASEA-RO) and Protease Inhibitor Cocktail tablet (ROCHE, cOmplete™ Sigma Aldrich cat. 11836170001). After 2h of incubation at 4°C, the soluble lysate was applied to a prepacked nickel column (HisTrap HP column, Cytiva Europe GmbH, France cat. GE17-5247-01) and fractionated on an AKTA pure system (Cytiva Europe GmbH, France) using a linear gradient from buffer A to buffer B (buffer A supplemented with 0.5 M imidazole) over 10 column volumes. A second step of purification was carried out using a Heparin affinity column (HiTrap Heparin HP Cytiva Europe GmbH, France cat. 17040703). The proteins were eluted using a buffer containing 50 mM Tris-HCl pH 7.5, 2 M NaCl, 1mM DL-Dithiothréitol (DTT), 0.1%, IGEPAL® CA-630 and 20% (v/v) glycerol.

### Microscale Thermophoresis (MST)

Microscale Thermophoresis (MST) utilizes an infrared (IR) laser to induce a precise temperature change in a sample, enabling the characterization and quantification of binding events through the monitoring of fluorescence changes. For this experiment, UPF1-HD, UPF1-CH-HD and UPF2 (770–1204) were labeled with the RED-tris-NTA 2nd Generation dye using the Monolith His-Tag Labeling Kit in accordance with the manufacturer’s instructions (Nano Temper Technologies, cat. MO-L018). Binding experiments were carried out using 200 nM of labeled protein and serial dilution of unlabeled ligand (as indicated in figures) in a Phosphate-based buffer as advised by the manufacture instructions. Samples were loaded into a premium-coated capillaries (NanoTemper, cat. MO-K02) and analyzed with NanoTemper Monolith NT.115 instrument set at 30°C with an excitation power of 60%. After data acquisition, the fluorescence traces were analyzed using software provided by the manufacturer. Binding curves are generated by plotting normalized fluorescence signals against the concentration of the binding partner. The binding affinity (Kd) and stoichiometry of the interaction were determined by fitting the data to appropriate binding models.

### Protein co-precipitation

Immunoprecipitations were carried out in 1.3×106 HEK 293T cells transfected with or without 2µg of the SARS-CoV-2 Np coding plasmid (addgene cat.141391). 48h after transfection, cells were washed two times with PBS and lysed with 1ml of lysis buffer (50 mM Tris–Cl, pH 8, 1% NP40, 0.5% sodium deoxycholate, 0.05% SDS, 0.1 mM EDTA, 150 mM NaCl, protease inhibitor (Roche)) for 30 min at 4°C. Half of the soluble fraction was incubated with 2µg of UPF1 antibody (bethyl) and the other half was incubated with 2µg of a rabbit pre-immune serum, overnight at 4°C. 10µl of mixed protein A and protein G magnetic beads were coated with 0.3% BSA and further added for 2h30 before extensive washings of 15min in lysis buffer supplemented with 0.1mg/ml RNAse A. Finally, beads were resuspended in laemlli buffer with loading dye to run western blots, as indicated.

The CBP pulldown procedure was conducted following the method outlined by (40,41) About 2 µg of each protein were combined in a binding buffer consisting of 20 mM Tris-HCl pH 7.5, 150 mM NaCl, 1 mM DTT, 1 mM MgCl2, 2mM CaCl2, and 10% (v/v) glycerol, and then incubated for 30 minutes at 37°C prior to resin mixing. Unbound proteins were removed by washing with binding buffer containing 200 mM NaCl. The proteins bound to the affinity beads were eluted using a buffer containing 20 mM EGTA, with the tubes shaken for 30 minutes at 30°C at 1400 rpm in a thermomixer. Eluates were separated by 12% SDS–PAGE and visualized through Coomassie staining.

### ATP hydrolysis

The steady-state experiments ATPase assays were carried out as described in (40). Before initiating the ATP assay, solutions containing either 5 µM of UPF1-HD or 5 µM UPF1-HD combined with 50 µM of Np, or 50 µM of Np alone, were incubated for 30 minutes at 37°C. Subsequently, 1 µl of each sample was incubated at 30°C in a 10-µl reaction mixture consisting of 20 mM MES pH 6.5, 100 mM potassium acetate, 1 mM DTT, 0.1 mM EDTA, 1 mM magnesium acetate, 1 mM zinc sulfate, 5% (v/v) glycerol, 2 µCi of [γ32P]-ATP (800 Ci/mmol, Revvity), 25 µM cold ATP, and 50 µM of polyU. At specified intervals, 2 µl reaction aliquots were withdrawn and quenched with 10 mM EDTA and 0.5% (v/v) SDS. Samples were then analyzed by phosphorimaging following thin-layer chromatography on polyethyleneimine cellulose plates (Merck) using 0.35 M potassium phosphate (pH 7.5) as the migration buffer.

### In vitro helicase assay

The 5’ single-strand overhang or blunt DNA duplexes were constructed using oligonucleotides as detailed in **Supplementary Figure 2A**. Initially, 50 pmol of radiolabeled FF2 was combined with 60 pmol of either FF1 or FF3 oligonucleotides in 100 µl of F100 buffer (20 mM MES pH 6.0, 100 mM potassium acetate, 1 mM DTT, 0.1 mM EDTA). The mixtures were heated to 95°C for 3 minutes and then gradually cooled to 20°C. Subsequently, DNA duplexes were purified on a native 6% (w/v) polyacrylamide gel (29:1) in 1x TBE buffer at 4°C. Single-run helicase assay was performed premixing the radiolabelled substrate (25 nM, final concentration) was mixed with an excess of UPF1, Np or preformed complexes (1 µM of UPF1) in F100 helicase buffer and incubated for 5 min at 30°C. The reaction was initiated by adding a mixture containing ATP and MgCl2 (1 mM, final concentrations), an excess of cold DNA strand (0.3 µM; to trap released DNA strands) and heparin (1 mg/ml; to trap free/released UPF1 and Np molecules). Reaction aliquots were withdrawn at various times, quenched with 150 mM sodium acetate, 10 mM EDTA, 0.5% (w/v) SDS, 25% (w/v) Ficoll-400, 0.05% (w/v) xylene cyanol, 0.05% (w/v) bromophenol blue and analysed by polyacrylamide gel electrophoresis and phosphorimaging on a Typhoon-Trio (GE Healthcare), as described in (41).

### Streptavidin displacement assay

The streptavidin displacement assay was conducted following the protocol outlined in (42). Using the helicase domain of UPF1, Np truncations, or preformed protein complexes as specified, a radiolabelled 3′-biotinylated 30-mer DNA substrate (**Supplementary Figure 2A**, from Sigma Aldrich) was bound to a Streptavidin monomer (Promega, cat. Z7041) in an F100 buffer.

For the displacement assay, 50 nM of DNA substrate was preincubated for 5 minutes at 30°C with 4 µM of UPF1-HD, Np, or preformed protein complexes in an F100 buffer. The enzymatic reaction was initiated by adding 2 mM ATP together with 4 µM of free biotin. At defined incubation times, 2 µl of reaction aliquots were mixed with 5 µl of quench buffer (150 mM NaAc, 10 mM EDTA, 0.5% (w/v) SDS, 25% (w/v) Ficoll-400, 0.05% (w/v) xylene cyanole, 0.05% (w/v) bromophenol blue) in preset tubes and immediately loaded onto the running gel. Samples were separated by an 8% (w/v) polyacrylamide gel containing 0.3% SDS and detected using a Typhoon 9400 phosphorimaging system and ImageQuant software (GE Healthcare).

### Electrophoretic mobility shift assay

The oligonucleotide FF2 (**Supplementary Figure2A**, Sigma Aldrich) and FF4 (**Supplementary Figure 2A**, Sigma Aldrich) were radiolabelled as indicated in (41) Protein complexes were prepared as indicated before then mixed with radiolabelled substrates (10 nM) with UPF1-HD (50 nM) with or without Np proteins in a buffer containing 20 mM MES pH 6.5, 150 mM potassium acetate, 2 mM DTT, 0.2 µg/µL BSA, and 6% (v/v) glycerol. The samples were incubated at 30°C for 20 min before being resolved by native 6.5% polyacrylamide (19:1) gel electrophoresis and analysed by phosphorimaging.

## Results

### Nucleocapsid protein of SARS-CoV-2 directly interacts with UPF1

Several lines of evidences indicated that Nucleocapsid protein (Np) from different CoVs is associated to cellular NMD inhibition via interaction with UPF1-containing RNP (18,35,36,38). Indeed, Np is associated with the incoming viral RNA genome (gRNA) and could represent the first line of defense against cellular RNA decay factors. To investigate the potential interaction between SARS-CoV-2 Np and endogenous UPF1, we initially transfected HEK 293T cells with a plasmid expressing Strep-tagged Np. Following RNAse A treatment to eliminate RNA-mediated interactions, we performed co-immunoprecipitations directed against endogenous UPF1 (**Figure 1A**). The effective interaction between UPF1 and Np was visualized by western blot using Anti-UPF1 and Anti-Strep-tag antibodies, respectively (**Figure 1A**).

**Figure 1.**
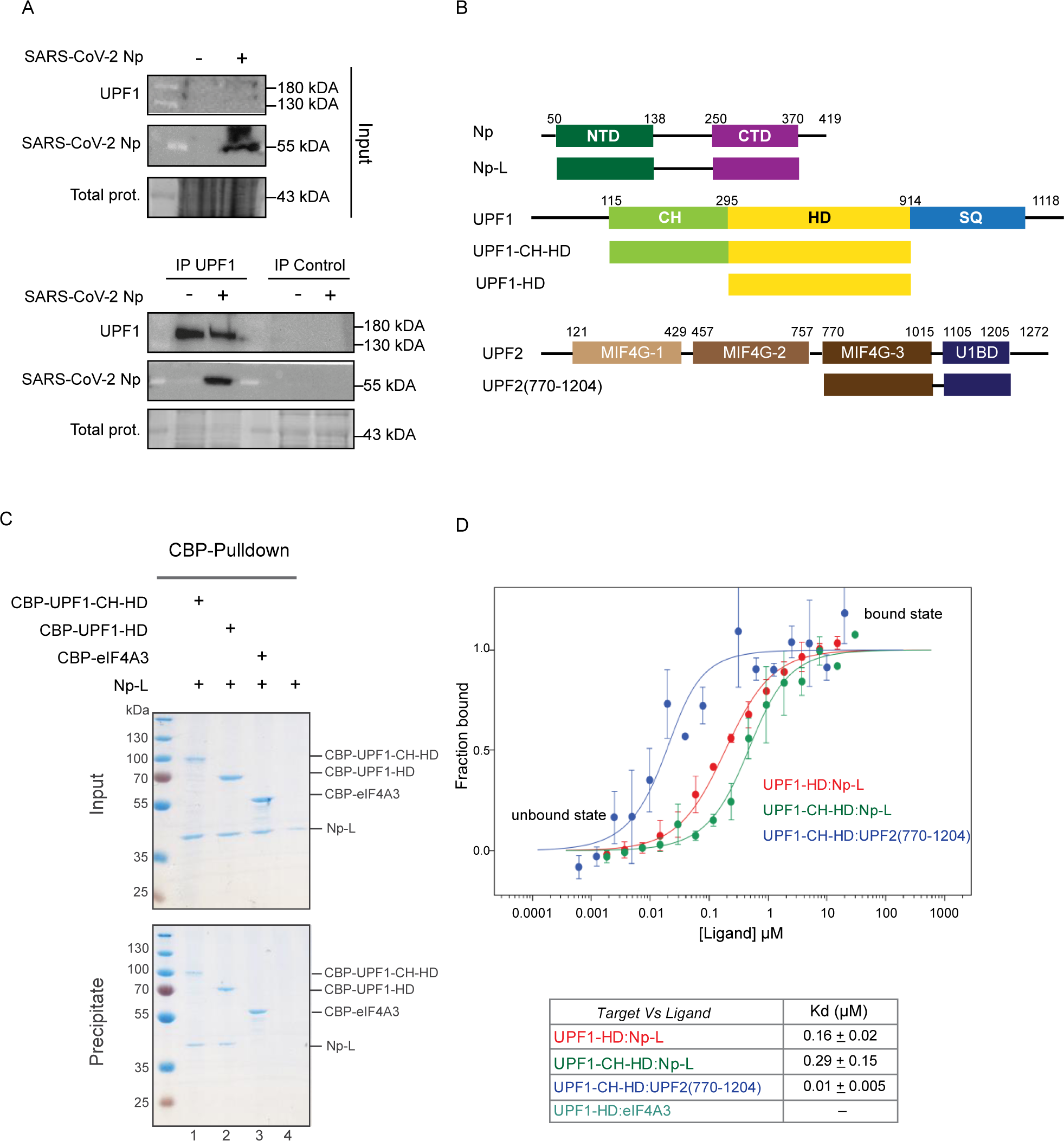
SARS-CoV-2 Np directly interacts with UPF1 in cellulo and in vitro. (A) Co-immunuprecipitation of endogenous UPF1 with SARS-CoV-2 Np in HEK 293T cells. Western blot of UPF1 and strep-tagged SARS-CoV-2 Np before (input, upper panel) and after immunoprecipitation (bottom panel) of endogenous UPF1. As a negative control, immunoprecipitation was also performed using a control serum. (B) Schematic diagram showing the SARS-CoV-2 Np, UPF1 and UPF2 domains used for this study. Structural domains are represented by rectangles and (predicted/confirmed) flexible regions by black lines. (C) Pulldown experiment using CBP-UPF1-CH-HD (lane 1), CBP-UPF1-HD (lane 2) or CBP-eIF4A3 (lane 3) as bait. After incubation, protein mixtures before (input 20% of total) or after precipitation (precipitate) were separated on a 12% SDS-PAGE gel and visualized by Coomassie staining. (D) MST quantification of UPF1 (target) –Np (ligand) interaction. The graph depicts the titration of Np-L or UPF2(770-1204) to a constant amount of UPF1-HD or UPF1-CH-HD fluorescently labeled. The binding curve represents the ligand concentration in a logarithmic scale, versus the proportion of the target that is bound by the ligand (fraction bound). The binding curve was fitted using a non-linear regression model to calculate the dissociation constant (Kd). Red and green curves are UPF1-HD and UPF1-CH-HD versus Np-L respectively; blue curve is UPF1-CH-HD versus UPF2(770-1204). Data points represent the mean values obtained from three independent experiments, with error bars indicating the standard deviation. At the bottom, the Kd values from MST experiments are summarized.

To further characterize the impact of Np on UPF1 activity, we performed an *in vitro* study of the SARS-CoV-2 Np-UPF1 interaction, using purified recombinant proteins and protein fragments. As in most nidoviruses, SARS-CoV-2 Np is composed of two structured domains: the N-terminal domain (Np-NTD) between amino acids [aa] 50 and 140 and the C-terminal domain (Np-CTD) between aa 250 and 370. Additionally, it contains two intrinsically disordered regions (IDRs) located at the N- and C-termini, as well as one Serine-Arginine (S/R) rich linker between the NTD and CTD (aa 140-250; **Figure 1B**). We used different truncations of Np: the Np-L (aa 50-370) depleted of the IDR extremities (**Figure 1B)**; the Np-NTD (aa 40-140), Np-NTD-SRD (aa 50-250) and Np-CTD (aa 250-330; **Supplementary Figure 1A**). Concerning UPF1 recombinant proteins, the boundaries between its three domains were defined according to previous structural and biochemical studies (40,43). We produced UPF1 containing the helicase domain (UPF1-HD aa 295-914) and a protein containing both, the HD and the N-terminal (CH) domain (UPF1-CH-HD; aa 115-914; **Figure 1B**) and a DEAD box helicase eIF4A3.

Calmodulin-binding peptide (CBP) and/or N- or C-terminal His-tagged proteins were purified as described in methods section. To assess the direct interaction of UPF1 and Np we incubated the two different UPF1 fragments with Np-L and performed CBP pulldown (**Figure 1C**). After extensive washes, input and eluted protein(s) were fractionated by SDS-PAGE and visualized by Coomassie staining. Np-L were co-precipitated when CBP-UPF1-CH-HD or CBP-UPF1-HD were used as bait (**Figure 1C**, lanes 1 and 2) and compared to CBP-eIF4A3 (**Figure 1C**, lane 3). Furthermore, microscale thermophoresis (MST) confirmed these interactions and determined a Kd of 160 nM for UPF1-HD:Np-L and 290 nM UPF1-CH-HD:Np-L (**Figure 1D** and **Supplementary Figure 1B**). MST assay using Np fragments only, indicated that the Np-CTD is domain interacts with better UPF1-HD than Np-NTD (**Supplementary Figure 1C**). As a control, we also used the UPF2 fragment containing the UPF1-binding region (aa 770-1204; **Figure 1B** and **Supplementary Figure 1B**) and found an expected very high affinity with UPF1-CH-HD resulting in a Kd of 10 nM (**Figure 1B** and **Supplementary Figure 1B**). Altogether, these results demonstrate the direct interaction of SARS-CoV-2 Np and UPF1

### Np inhibits the unwinding activity of UPF1 and stabilizes a NMD-prone RNA substrate

UPF1 utilizes the energy derived from ATP hydrolysis to translocate on DNA or RNA substrates in a 5′-3′ direction. This translocation enables UPF1 to unwind nucleic acid (NA) duplexes or displace proteins located on the substrate (40,42,43). To assess whether SARS-CoV-2 Np impacts UPF1 activity, we initially monitored the enzyme’s ability to unwind a DNA duplex. We chose DNA because it has been observed that the unphosphorylated Np forms biomolecular condensates in the presence of RNA ((44–51). The DNA substrate was composed of a 12-nucleotide (nt) 5’ single-stranded DNA (ssDNA) followed by a 21-base-pair (bp) double-stranded DNA (dsDNA) region (**Figure 2A and Supplementary Figure 2A**). The substrate was first incubated with UPF1-HD, which exclusively binds to the ssDNA portion. Subsequently, the unwinding reaction was triggered by ATP addition and the unwinding reaction was monitored under a pre-steady state regimen (see methods) by tracking the radiolabeled bottom-strand DNA oligonucleotide (**Figure 2A**; (40,41)). Under these conditions, only the first round of unwinding was monitored during the reaction (single-run conditions). Unwinding efficiencies were determined based on the proportions of 32P-labeled single-stranded DNA accumulated over time (**Figure 2A** and **2B**).

**Figure 2.**
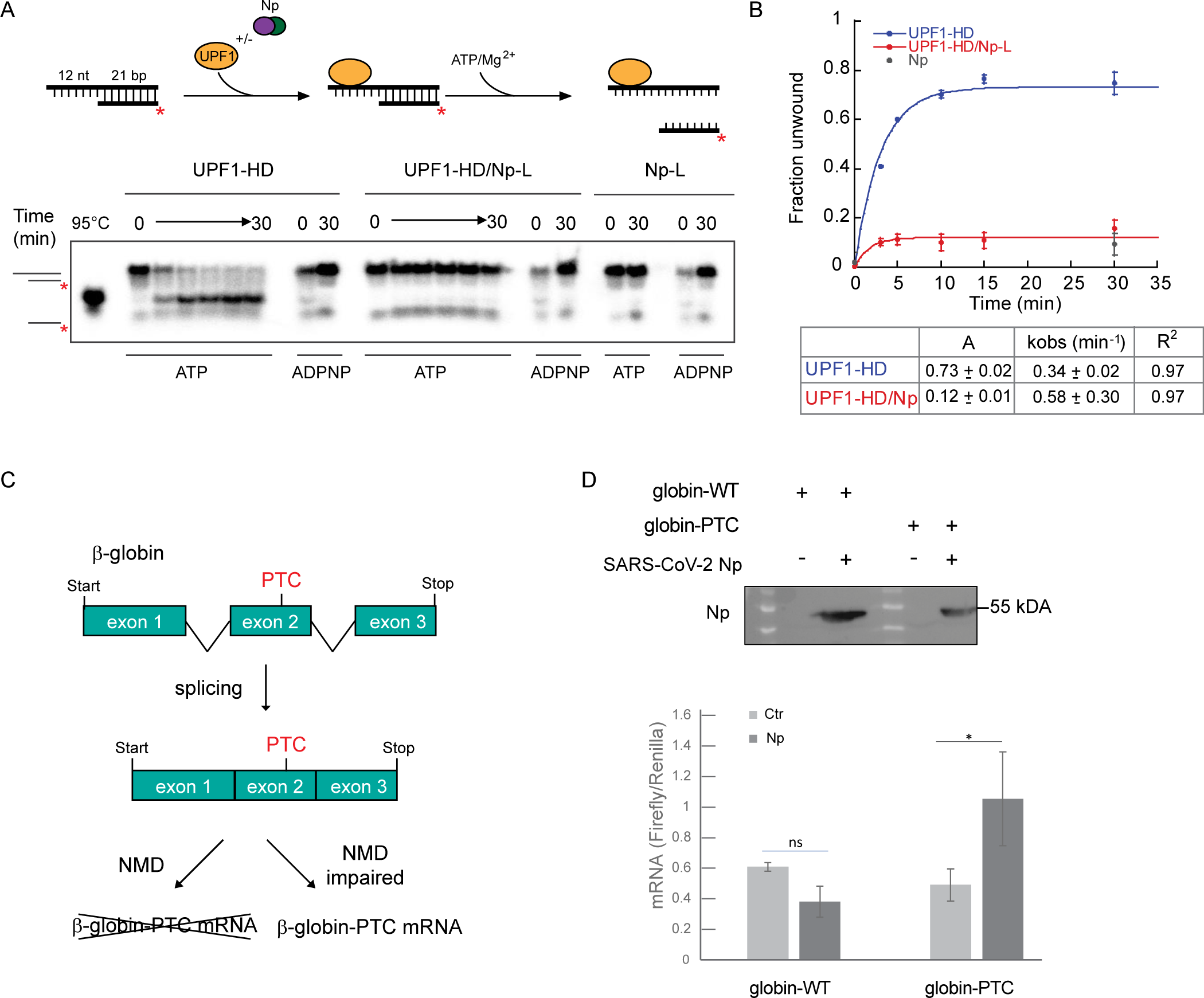
SARS-CoV-2 Np inhibits UPF1 unwinding activity and cellular NMD. (A) A representative gel showing ATP-induced unwinding of the dsDNA substrate with 5’ overhangs of 12 nucleotides by UPF1-HD, the UPF1-HD:Np-L preformed complex, and Np-L alone. The substrate and the experimental workflow are depicted at the top of the gel. In the control lanes, ATP was replaced by ADPNP or samples were heat-denatured (95°C). (B) Graph showing the fraction of DNA oligonucleotide released over time. Data points derived from four independent experiments were fitted to the pseudo-first order equation y=A[1-e(-kt)], where A and k represent, respectively, the amplitude and the rate constant of the unwinding reaction. (C) Schematic representation of NMD assay to assess the stability of mRNA expressed from a β-globin reporter minigene that was either WT (Gl-WT) or with a PTC in the second exon (Gl-PTC). (D) RNA decay assay was performed in cells overexpressing or not the SARS-CoV-2 Np. In the upper panel, the western blot shows the overexpression level of SARS-CoV-2 Np. NMD efficiency was determined by the quantitative mRNA detection of NMD reporters. Relative levels of Gl-WT and Gl-PTC mRNAs were determined by RT-qPCR. The statistical significance was supported by a T test with three independent biological replicates.

Remarkably, the unwinding activity of UPF1-HD is strongly affected by the presence of the Np-L protein (**Figure 2A** and **2B**). Indeed, the amplitude of the reaction with UPF1-HD:Np-L is reduced by approximatively 6-fold when compared to that obtained with Upf1-HD alone (**Figure 2B**). The Np-NTD and Np-CTD truncations presented a half level of UPF1-HD unwinding inhibition compared to the Np-L (**Supplementary Figure 2B**). As expected, when ATP is replaced by non-hydrolysable ATP analog in the reaction, ADPNP, the fraction of duplex unwound over time is negligible (**Figure 2A**). These data clearly demonstrate that only the Np-L fully inhibits the helicase activity of UPF1.

It is possible that this inhibition of UPF1 enzymatic activity may result in an inhibition of NMD, as already observed for MHV coronavirus (Wada et al 2018). Thus we performed a NMD assay in HeLa cells expressing SARS-CoV-2 Np. In this test, we analyzed the level of mRNAs transcribed from transfected β-globin minigenes, WT (Gl-WT) or with a premature translation termination codon (Gl-PTC) (**Figure 2C**). Both transcripts undergo splicing, and while the Gl-PTC reporter is targeted by NMD and degraded, the Gl-WT is not (**Figure 2C**). The quantitative reverse transcription–polymerase chain reaction (RT-qPCR) analysis revealed that Gl-PTC transcript is significantly stabilized by the presence of Np compared to Gl-WT mRNA in the same conditions (**Figure 2D**). This result indicates that Np inhibits cellular NMD.

### The Np physically inhibits the progression of UPF1 on the substrate

The physicochemical properties of SARS-CoV-2 Np allow for its nonspecific binding to nucleic acids, including both ssDNA and dsDNA, *in vitro* (52–55). This raises the issue that Np could affect UPF1 through an indirect mechanism, primarily acting on the nucleic acid substrate. To understand how Np affects UPF1’s unwinding activity, we studied the enzymatic properties associated with its helicase. As mentioned, unwinding is fueled by ATP hydrolysis, so we investigated the effect of Np on this process. Indeed, the ATP hydrolysis activity of UPF1-HD is identical whether Np-L is present or not suggesting rather an indirect mechanism. (**Figure 3A**).

**Figure 3.**
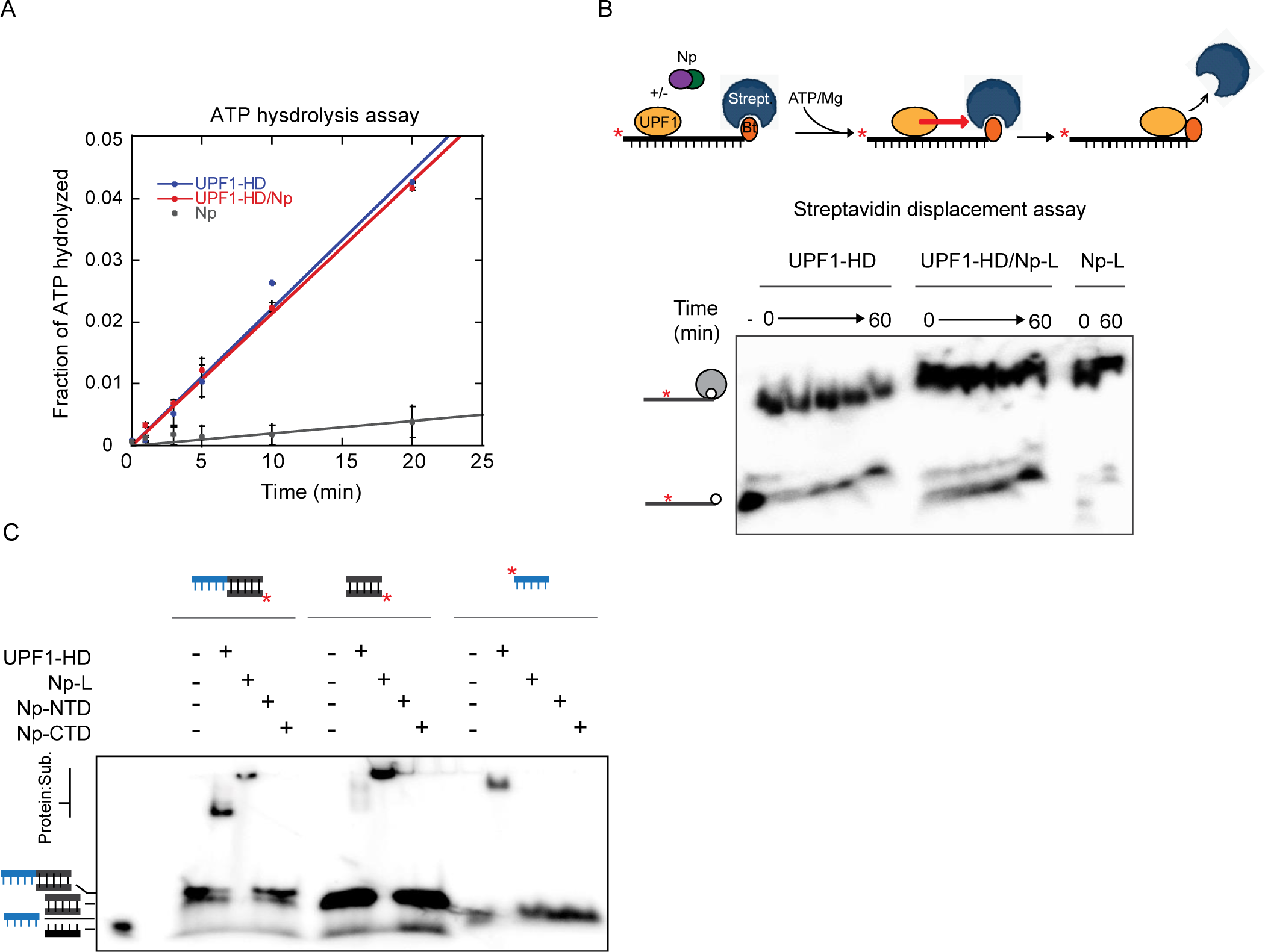
SARS-CoV-2 Np does not affect the unwinding-associated functions of UPF1. (A) Graph showing the percentage of ATP hydrolysed as a function of time by the UPF1-HD, UPF1-HD:Np-L preformed complex, and Np-L alone under conditions of steady-state ATPase turnover (see ‘Materials and Methods’ section). (B) Time course of an active displacement of streptavidin from a 3’ biotinylated DNA by UPF1-HD, the UPF1-HD:Np-L preformed complex, and Np-L alone. The pipeline of the experiment is showed at the top of the gel. At various times (see Methods section) aliquots from the helicase reaction were mixed in a quench buffer and loaded immediately on 8% polyacrylamide gel containing 0.3% of SDS to selectively denature protein/s:DNA complexes without altering biotin-streptavidin interaction. (C) Representative native 6.5% polyacrylamide gels illustrating the interaction of UPF1-HD and Np-L, Np-NTD and Np-CTD proteins with different DNA substrates depicted at the top of the gel. The radiolabeled 5’ extremity is showed by a red star, the overhang DNA is in blue and the dsDNA in black. The substrates were incubated with the same concentrations of proteins used in helicase or streptavidin displacement assays.

Then, we focused on the nucleoprotein remodeling activity of UPF1. UPF1 is able to displace streptavidin bound to a biotinylated RNA substrate or single-stranded DNA binding proteins coating the DNA by translocation (42,56). To measure this activity we conducted a streptavidin displacement assay: briefly, UPF1-HD and Np-L were pre-incubated at 37°C then a radiolabeled substrate is added as indicated in **Figure 3B** (upper scheme), before the reaction was initiated by adding ATP, magnesium, and an excess of biotin to trap free streptavidin molecules. Time-course aliquots were then analyzed on a native polyacrylamide gel to separate free DNA from streptavidin-bound DNAs. Remarkably, both UPF1-HD and the UPF1-HD:Np-L complex efficiently displaced the streptavidin from the substrate indicating that the translocation ability of UPF1 is not affected by the presence of SARS-CoV-2 Np (**Figure 3D**). The same efficient streptavidin displacement was observed using UPF1-HD:Np-NTD and UPF1-HD:Np-CTD protein complexes (**Supplementary Figure 2C**).

One possible explanation for the inhibition of helicase activity without alteration of ATP hydrolysis or translocation is that Np physically binds to the DNA substrate downstream of UPF1 binding site, thereby obstructing the progression of the enzyme. To test this hypothesis, we conducted an electrophoretic mobility shift assay (EMSA) under the same conditions as the enzymatic tests, using isolated portions of the DNA duplex substrate (Supplementary Figure 2A, **Figure 3C**). We used the entire DNA duplex, the 12-nts 5’ ssDNA overhang and the blunt dsDNA region. Each radiolabeled substrate was incubated with the indicated protein truncations, at the same concentration and under the same conditions used for the enzymatic tests. Our observations revealed that while UPF1-HD interacted with ssDNA only as anticipated, Np-L interacted with the dsDNA region of the substrate in our conditions (**Figure 3C**). Neither the Np-NTD nor the Np-CTD protein fragments exhibited stable interactions with either ssDNA or dsDNA (**Figure 3C**). This outcome corroborated the hypothesis that SARS-CoV-2 Np hampers the unwinding activity of UPF1 by binding to the structured region of the substrate.

### SARS-CoV-2 Np binds also with UPF2 preventing its interaction with UPF1

The physical block provided by Np for UPF1 progression and unwinding supports a model in which Np protects the incoming SARS-CoV-2 genome by binding to both structured and linear portions of the viral RNA (48). This protection might explain how Np safeguards the viral genome from UPF1, preventing its degradation by the cellular NMD pathway, but does not account for the effects observed on cellular NMD target RNAs (**Figure 2C**).

This NMD inhibition mediated by Np should occur through a mechanism distinct from physical RNA protection. Thus we tested the hypothesis that the activation step of UPF1 enzymatic exerted by UPF2 during NMD could be affected by the direct interaction of Np to UPF1. We performed a CBP pulldown of UPF1-CH-HD, UPF2(770–1204) fragment and SARS-CoV-2 Np-L (**Figure 4A**). As expected, UPF2(770–1204) or Np-L were co-precipitated when CBP-UPF1-CH-HD was used as bait (**Figure 4A**; lanes 1 and 2) whereas only Np-L was co-precipitated by UPF1-CH-HD when present together with UPF2(770–1204) in the reaction mixture (**Figure 4A**; lane3). This result was quite surprising, as the affinity measured for the UPF1-CH-HD:UPF2(770–1204) interaction is about 30 times higher than for the UPF1-CH-HD:Np-L (**Figure 1D** and **Supplementary Figure 1B**). Thus we examined the possibility that Np-L interacts also with UPF2. Surprisingly, using MST, we observed an interaction between UPF2(770–1204) and SARS-CoV-2 Np with an affinity comparable to that of UPF1-CH-HD and UPF2(770–1204) under these conditions. (**Figure 4B**). However, we observed differences in the Kd values obtained when measuring the affinity of UPF1-CH-HD and UPF2(770–1204) depending on which protein is labeled for the experiment. These differences are likely attributable to the presence of the fluorophore attached to UPF2, which can cause steric hindrance and decrease its affinity for UPF1-CH-HD.

**Figure 4:**
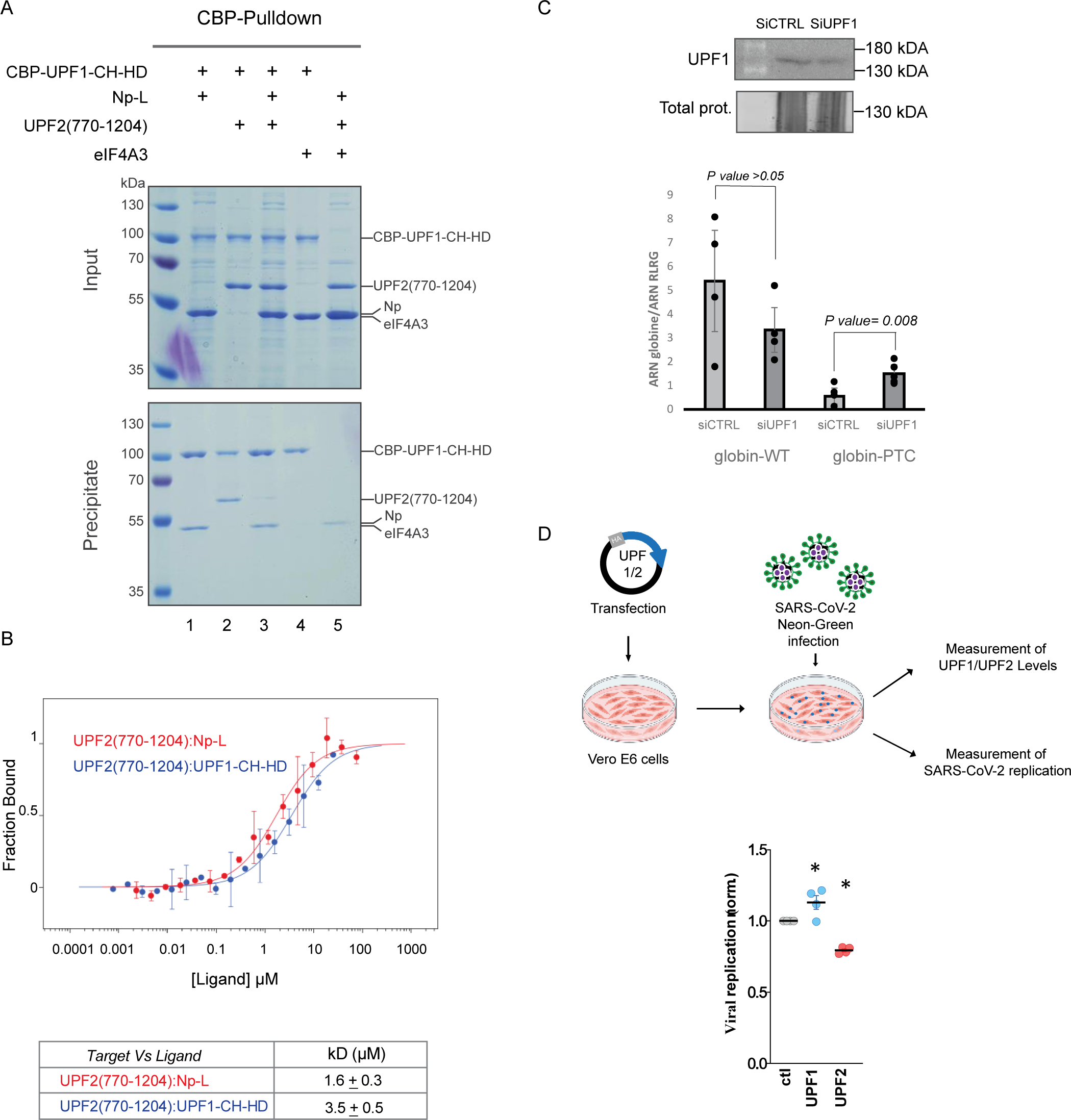
SARS-CoV-2 Np interacts with UPF1 and UPF2 preventing their interaction. (A) Pulldown experiment using CBP-UPF1-CH-HD (lanes 1 to 4) as bait and UPF2(770-1204), SARS-CoV-2 Np and eIF4A3 as preys. After incubation, protein mixtures before (input 20% of total) or after precipitation (precipitate) were separated on a 12% SDS-PAGE gel and visualized by coomassie staining. (B) MST quantification of UPF2 (target) – Np or -UPF1 (ligands) interactions. The graph depicts the titration of Np-L or UPF1-CH-HD to a constant amount of UPF2(770-1204) fluorescently labeled. The binding curve represents the ligand concentration in a logarithmic scale, versus the proportion of the target that is bound by the ligand (fraction bound). The binding curve was fitted using a non-linear regression model to calculate the dissociation constant (Kd). Red and blue curves are UPF2(770-1204) versus Np-L or UPF1-CH-HD respectively. Data points represent the mean values obtained from three independent experiments, with error bars indicating the standard deviation. At the bottom, the Kd values from MST experiments are summarized. (C) Western blot analysis of UPF1 protein expression after 24 h transfection of Vero E6 cells with siRNAs targeting UPF1 compared to a control siRNA (upper panel). Quantification of Gl-WT or Gl-PTC mRNAs from cells treated with the indicated siRNA (bottom panel). The statistical significance was supported by a T test with three independent biological replicates. (D) Vero E6 were transfected with DNA coding for HA-UPF1, HA-UPF2, or a control DNA. Twenty eight hours later they were infected with MOIs comprised between 0.01 and 0.01 with a Neon-Green bearing replicative-competent SARS-CoV-2 virus. The extent of viral replication and the overexpression of UPF1/2 were determined by flow cytometry eighteen hours post viral challenge by flow cytometry (upper panel). The graph presents averages and SEM from 4 independent experiments. *, p<0.05, following an ordinary one-way Anova test between the indicated conditions (bottom panel).

In conclusion, the inhibition of NMD can be explained by the competition exerted by Np, which prevents the interaction between UPF1 and UPF2 and the subsequent activation of UPF1.

### SARS-CoV-2 replication is modulated by ectopic expression of NMD factors

UPF1 and UPF2 are the key players in NMD. According to the results obtained above, we hypothesized that their overexpression in the context of SARS-CoV-2 replication should lead to a decrease in viral replication. To test this hypothesis, we first monitored the activity of UPF1 in the African Green monkey kidney cell line Vero E6, a suitable cellular model for studying SARS-CoV-2 replication using reporters of NMD. To this end, Vero E6 were treated with siRNA either control or targeting UPF1 and we analyzed the level of mRNAs transcribed from transfected β-globin minigenes, WT (Gl-WT) or with a premature translation termination codon (Gl-PTC) as described before. As expected, RT-qPCR analysis revealed that Gl-PTC is less stable than Gl-WT in control conditions. Moreover, under UPF1-depletion condition, it showed a specific three-time accumulation of Gl-PTC reporter transcript, confirming the degradation of Gl-PTC reporter transcript by the NMD pathway and the correct activity of UPF1 in Vero E6 cells (**Figure 4C**).

Following these results, we ectopically expressed UPF1 and UPF2 in VeroE6 cells prior to viral challenge with a SARS-CoV-2 strain bearing a NeonGreen reporter. The extent of viral replication was then assessed at eighteen hours post infection by flow cytometry (**Supplementary Figure 3**). This relatively early time point was chosen to minimize unwanted side effects that may be due to UPF1/2 overexpression on the cell. Under these conditions, the replication of SARS-CoV-2 was enhanced, albeit slightly, by ectopic expression of UPF1 and in contrast, it was inhibited upon overexpression of UPF2 (**Figure 4D**). Overall, these data suggest that UPF proteins modulate SARS-CoV-2 replication.

## Discussion

In the study of the ongoing arms race between viruses and hosts, increasing evidence shows that NMD is indeed a genuine defense mechanism against several positive-strand RNA viruses. Here, we showed the direct interaction of SARS-CoV-2 Np with UPF1 and UPF2 and demonstrated its consequences on inhibiting the NMD process. We observed a complex multi-level mechanism of UPF1 enzymatic regulation exerted by Np. The first level of regulation is exerted against the unwinding activity of UPF1. Although its ATP hydrolysis and translocation/remodeling activities remain unaffected by Np, the unwinding is strongly inhibited.

To shed light on this surprising effect, we performed a combination of bulk enzymatic assays using DNA substrates. The choice of DNA was motivated by the fact that UPF1 is not specific to a single nucleic acid *in vitro* and acts in several RNA- and DNA-related pathways in the cell (reviewed in (57) and (58)). Moreover, the Np of SARS-CoV-2 has also been shown to promiscuously bind ss- and ds-nucleic acids *in vitro* (52–55) but produces condensates only in the presence of ss- or dsRNA, which could bias the enzymatic study (44,45,47–51,59).

Using this DNA setup, we observed that UPF1 unwinding inhibition primarily occurs through the physical binding of Np to the double-stranded portion of the DNA substrate, thereby impeding the progression of UPF1. Even though Np has been shown to bind ssDNA in single-molecule studies (52), under our conditions, Np does not. However, the preference of Np for ssRNA sites flanked by stable stem-loops has been described (48,59), suggesting that Np could bind ssDNA when followed by a double-stranded region. Interestingly, until now, neither the presence of an avid ssDNA binder such as Gp32-B nor the presence of a biotin/streptavidin complex on ssRNA or ssDNA substrates has been able to impede the enzyme’s progression (42). Notably, this loss of UPF1 accessibility to double-stranded substrate is consistent with the idea that Np, by tightly condensing the viral genome, serves as the first line of defense against cellular restriction factors.

The second level of regulation involves the direct interaction of SARS-CoV-2 Np with UPF1. This finding aligns with a proteomic study that identified the Np of SARS-CoV-2 as a UPF1 interactor (36), as well as with previous studies on other coronaviruses, where Np has been shown to inhibit NMD by interacting with UPF1-containing RNP complexes (18,35). In our experiments, Np binds to both UPF1 and UPF2 with micromolar or sub-micromolar dissociation constants (Kd), indicating a strong binding affinity. This interaction prevents UPF1 and UPF2 from interacting: an essential step for the progression of NMD and RNA decay.

In this work, we also showed that UPF1 overexpression in SARS-CoV-2 infected cells slightly increased viral replication, while overexpression of UPF2 decreased it. While the observation that an increased expression of UPF2 can inhibit viral replication supports the hypothesis that SARS-CoV-2 inhibits NMD, the finding that UPF1 overexpression increases it, is at first sight at odd with this hypothesis. At present, the reasons for this are unclear and it remains possible that despite the fact that they interact with each other, UPF1 and UPF2 may also play distinct additional roles in RNA metabolism.In support of this hypothesis, UPF1 overexpression enhanced viral RNA stability and translation of HIV-1, thereby promoting the production of viral proteins and the assembly of new virions (14,31,32), while UPF2 inhibits HIV-1 viral protein expression by promoting viral RNA decay (3,4,33,51).

In summary, our study elucidates a multifaceted strategy employed by SARS-CoV-2 to evade NMD. By directly interacting with UPF1, the Np protein not only inhibits the helicase activity of UPF1 but also disrupts the UPF1-UPF2 interaction and usurps UPF1 for the advantage of viral replication (**Figure 5**). This mechanism likely enhances the stability of viral RNAs, contributing to efficient viral replication. These findings highlight the importance of UPF1 in SARS-CoV-2 replication and suggest potential therapeutic targets within the NMD pathway to combat SARS-CoV-2 infection. Further research is needed to fully understand the broader implications of these interactions and to explore potential interventions that could disrupt the virus’s ability to manipulate host RNA decay pathways.

**Figure 5:**
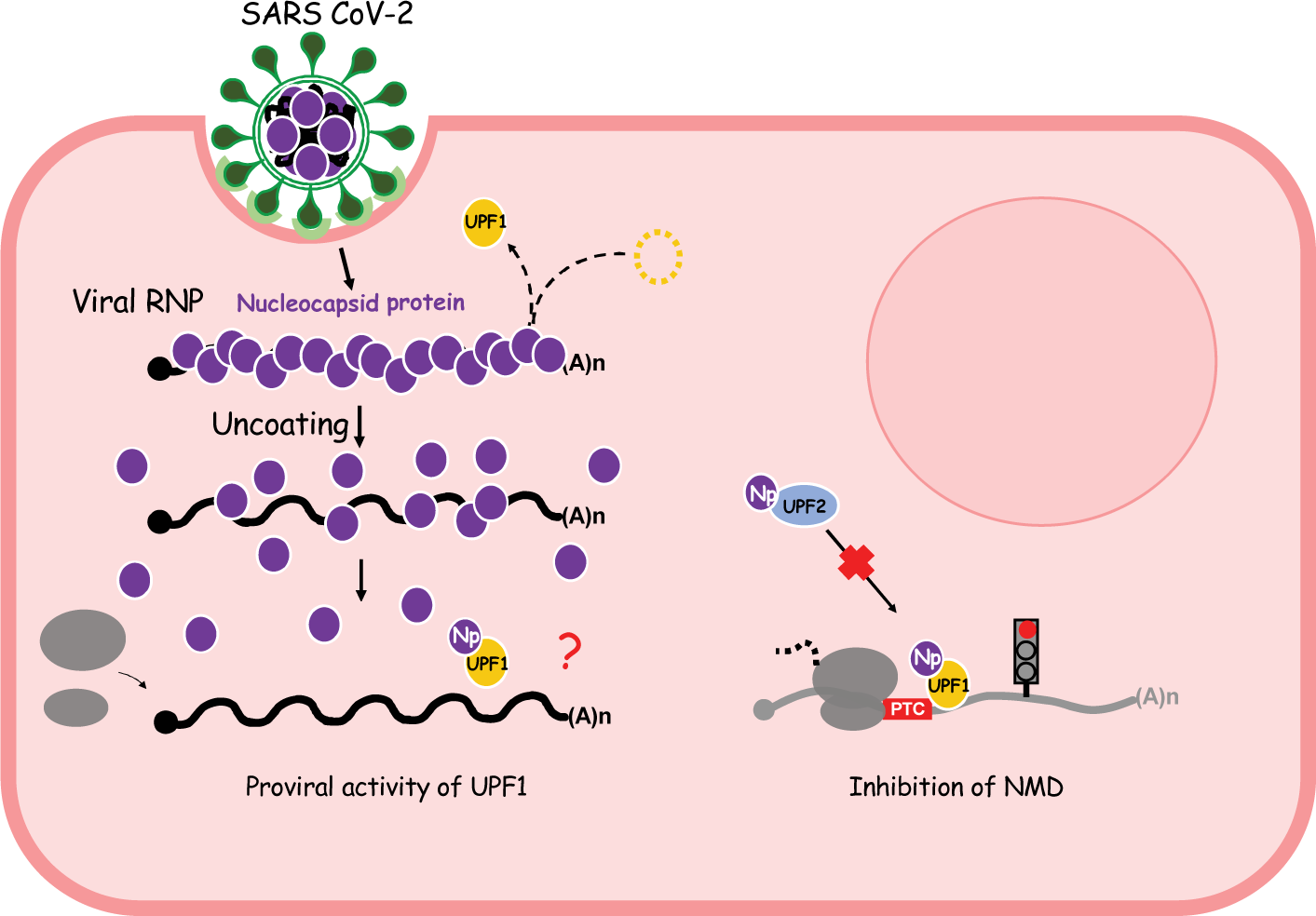
proposed model.

## Acknowledgements

We thank Dr. Christine Roden from the University of North Carolina at Chapel Hill, USA, for kindly providing us with the expression plasmid for SARS-CoV-2 Np. We acknowledge the support of the Protein Science Facility (PSF) of SFR Biosciences (UAR3444/CNRS, US8/Inserm, ENS de Lyon, UCBL), especially Virginie Gueguen-Chaignon and Céline Freton for their assistance, and Frédéric Galisson for his help with radioactive sources.

## Author contributions

V.M., A.C. and F.F. conceptualized the study and conceived and designed experiments. V.N., F.G.B., C.R., P.D.C. acquired and analyzed the majority of the *in vitro* biochemical experiments. M.M.H., S.D., A.R. performed cellular and viral experiments. F.F. wrote the manuscript and all other authors edited it.

## Supplementary material

**Supplementary Figure 1:**
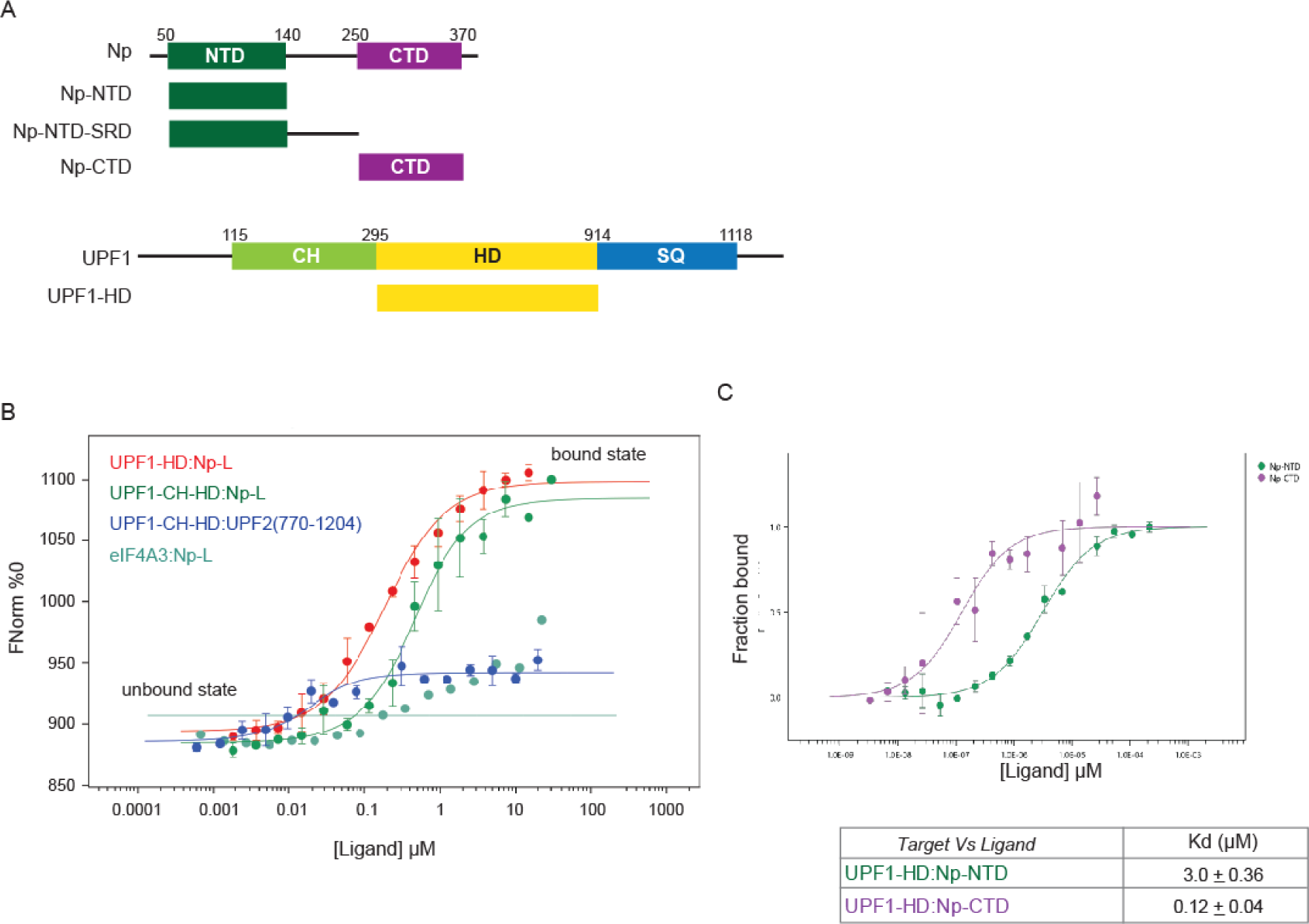
(A) Schematic diagram showing the SARS-CoV-2 Np truncations, UPF1 domains used for this study. Structural domains are represented by rectangles and (predicted/confirmed) flexible regions by black lines. (B) MST quantification of UPF1 (target) –Np (ligand) interaction. The graph depicts the experiment of Figure 2D with binding curve representing the ligand concentration in a logarithmic scale, versus the normalized fluorescence (Fnorm in ‰). Red and green curves are UPF1-HD and UPF1-CH-HD versus Np-L respectively; blue curve is UPF1-CH-HD versus UPF2(770-1204) and blue/green curve is the RNA helicase eIF4A3 versus Np-L. (C) MST quantification of UPF1 (target) –Np-NTD or Np-CTD (ligands) interaction. The graph depicts the titration of Np-NTD (green) and Np-CTD (purple) to a constant amount of UPF1-HD fluorescently labeled. The binding curve represents the ligand concentration in a logarithmic scale, versus the proportion of the target that is bound by the ligand (fraction bound). The binding curve was fitted using a non-linear regression model to calculate the dissociation constant (Kd). Data points represent the mean values obtained from three independent experiments, with error bars indicating the standard deviation. At the bottom, the Kd values from MST experiments are summarized.

**Supplementary Figure 2:**
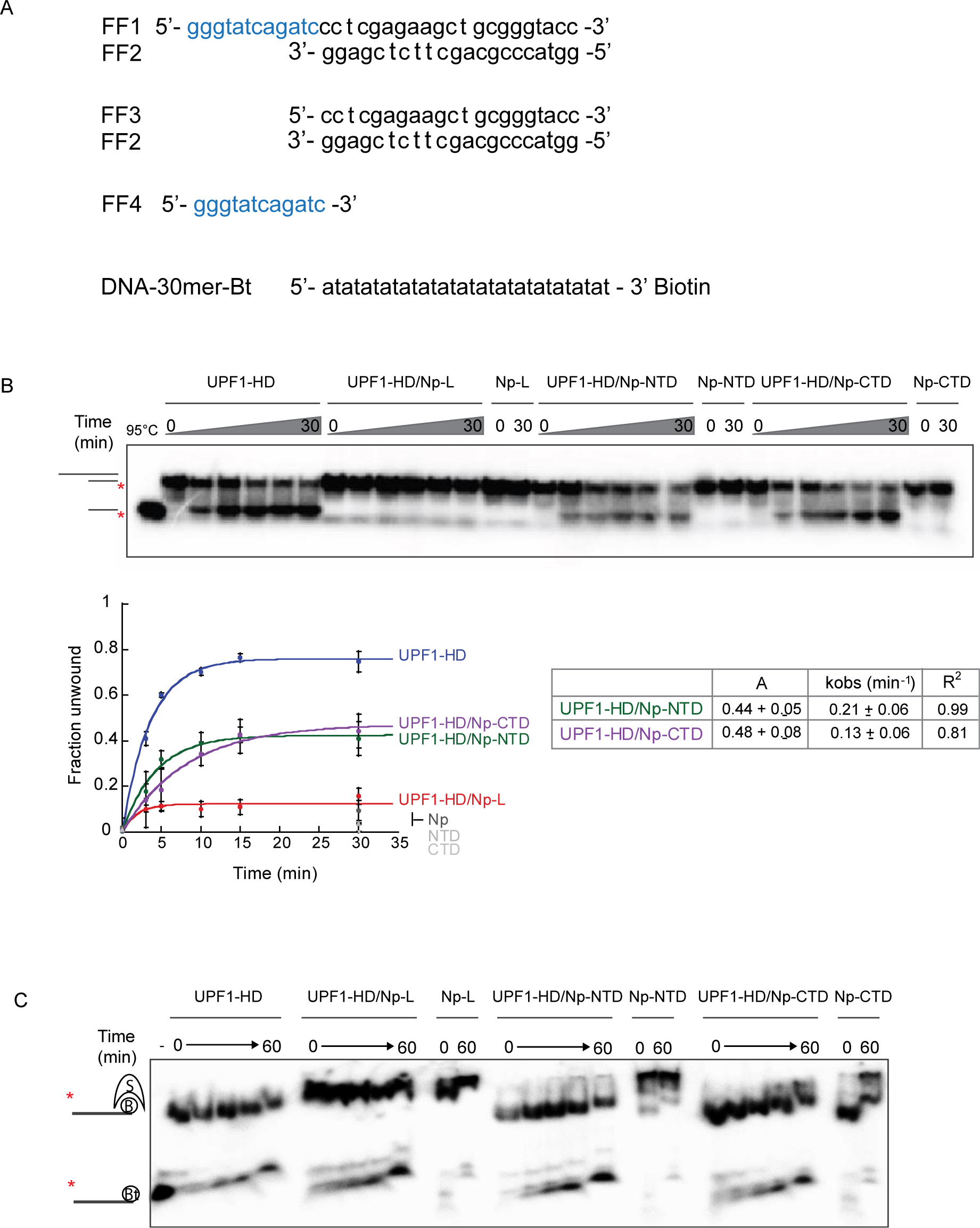
SARS-CoV-2 Np-NTD and CTD partially inhibits UPF1 unwinding activity without affecting RNP remodeling. (A) A representative gel showing ATP-induced unwinding of the dsDNA substrate with 5’ overhangs of 12 nucleotides by UPF1-HD, the UPF1-HD:Np-L; UPF1-HD:Np-NTD; UPF1-HD:Np-CTD preformed complexes, and Np truncations. In the control lanes, ATP was replaced by ADPNP or samples were heat-denatured (95°C). At the bottom the graph shows the fraction of DNA oligonucleotide released over time. Data points derived from four independent experiments were fitted to the pseudo-first order equation y=A[1-e(-kt)], where A and k represent, respectively, the amplitude and the rate constant of the unwinding reaction. (B) Time course of an active displacement of streptavidin from a 3’ biotinylated DNA by UPF1-HD, the UPF1-HD:Np-L; UPF1-HD:Np-NTD; UPF1-HD:Np-CTD preformed complexes, and Np truncations. At various times (see Methods section) aliquots from the helicase reaction were mixed in a quench buffer and loaded immediately on 8% polyacrylamide gel containing 0.3% of SDS to selectively denature protein/s:DNA complexes without altering biotin-streptavidin interaction.

**Supplementary Figure 3:**
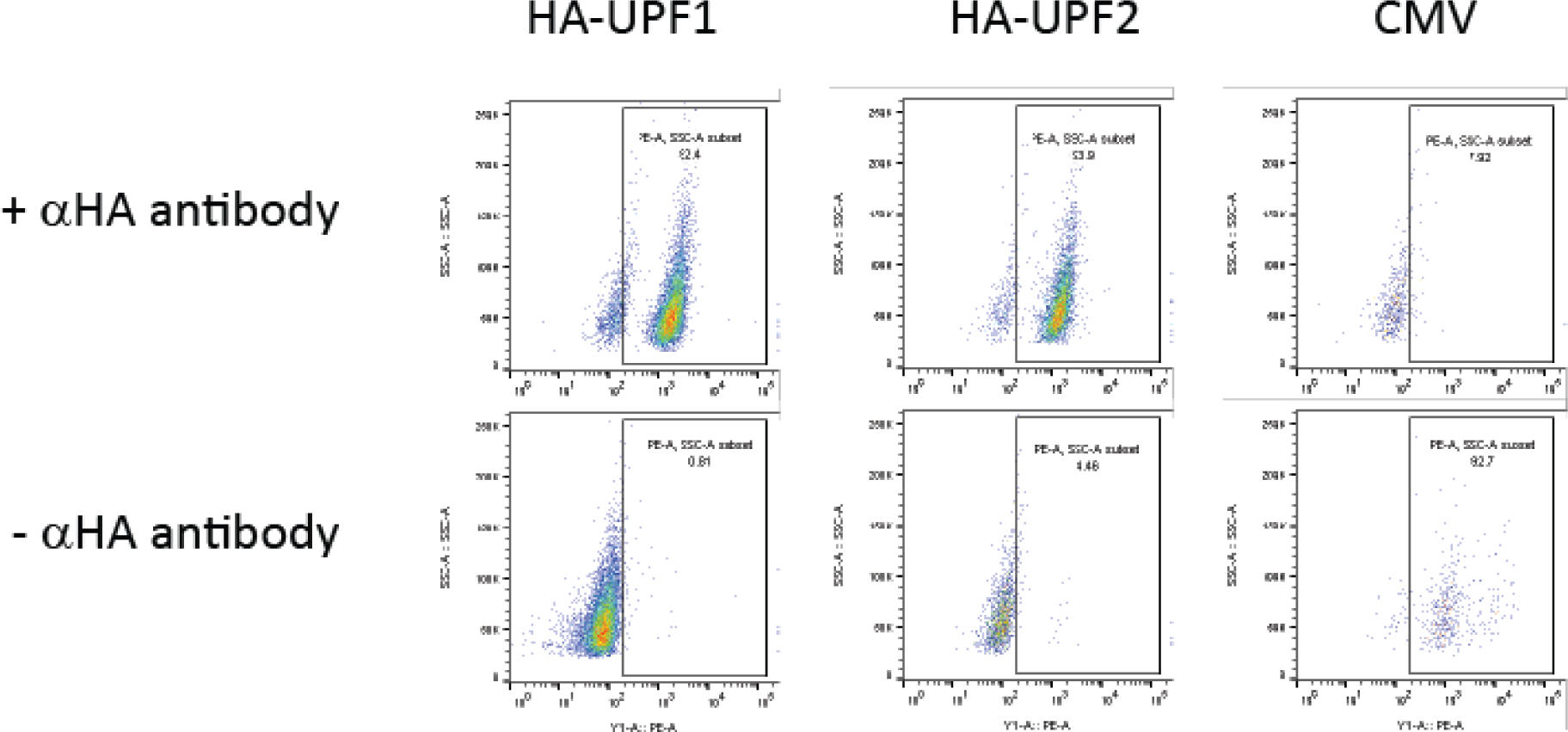
Validation of HA UPF1 and HA UPF2 Overexpression (relative to Figure 4D) by FACS using a primary antibody targeting the HA tag.

